# Eukaryotic Plankton Community Assembly and Influencing Factors Between Continental Shelf and Slope Sites in the Northern South China Sea

**DOI:** 10.1101/2022.02.09.479823

**Authors:** Tangcheng Li, Guilin Liu, Huatao Yuan, Jianwei Chen, Xin Lin, Hongfei Li, Liying Yu, Cong Wang, Ling Li, Yunyun Zhuang, Senjie Lin

## Abstract

Eukaryotic plankton are pivotal members of marine ecosystems playing crucial roles in food webs and biogeochemical cycles. However, understanding the patterns and drivers of their community assembly remains a grand challenge. A study was conducted in the northern South China Sea (SCS) to address this issue. Here, 49 samples were collected and size-fractionated from discrete depths at continental shelf and continental slope in the northern South China Sea over a diel cycle. From high throughput sequencing of the 18S rDNA gene V4 region, 2,463 operational taxonomic units (OTUs) were retrieved. Alveolata and Opisthokonta overwhelmingly dominated the assemblages in relative abundance (44.76%, 31.08%) and species richness (59%, 12%). Biodiversity was higher in the slope than the shelf and increased with depth. Temperature and salinity were found as the major deterministic drivers of taxon composition. Community structure was influenced by multiple factors in the importance order of: environmental factors (temperature + salinity) > spatial factor > water depth > sampling time. Furthermore, the neutral model explained more variations in the smaller-sized (0.22-3μm) community (24%) than larger-sized (3-200 μm) community (16%) but generally explained less variations than did deterministic processes. Additionally, our data indicated that the larger plankton might be more environmentally filtered and less plastic whereas the smaller plankton had stronger dispersal ability. This study provides novel insights into differential contributions of the deterministic process and stochastic process and complexities of assembly mechanisms in shaping the community assembly of micro-nano and pico-eukaryotic biospheres in a subtropical ocean.

**Importance:** Eukaryotic plankton are essential to the biogeochemical processes in the ocean. Understanding the plankton community and influence factors is rapidly increasing with the advances in high-throughput sequencing in recent years, especially the microbe. Our study documented the biodiversity and drivers of community assembly of eukaryotic plankton in the subtropic northern SCS. We found the community structure was influenced by multiple factors and the deterministic process was more important than the stochastic process in the assembly of the eukaryotic plankton communities. Temperture is the most important factors in deterministic factors rather the nutrients which were previously thought important in shaping the eukaryotic plankton community in this oceanic province. Furthermore, we found smaller-sized miceukaryotic plankton are less environment filtered and more plastic than the larger-sized plankton. This insight revealed the differential contributions of the deterministic process and stochastic process and complexities of assembly mechanisms in shaping the community of eukaryotic biospheres.

## Introduction

Microeukaryotes are highly diverse and play fundamental roles as producers, consumers, and trophic links in aquatic food webs [1, 2]. They are essential to the biogeochemical processes in the ocean, with phytoplankton fixing CO2 and other elements into organic matter that lead to the export of carbon to the deep ocean [1]. Our understanding of the biodiversity and the structure of marine microeukaryotic communities and influencing factors is rapidly increasing with the advances in high-throughput sequencing [3–9]. Most notably, The TARA Oceans expedition captured 150,000 operational taxonomic units (OTUs) in the sunlit ocean across the globe, remarkably enriching the marine life inventory [5]. However, compared to bacteria, marine eukaryotic plankton communities are still much less explored for natural biodiversity [10].

The size of phytoplankton has profound impacts on marine ecology and biogeochemical cycling [11]. Due to a smaller thickness of the diffusion boundary layer and a larger surface-area-to-volume ratio, smaller-sized phytoplankton has a competitive advantage over larger-sized phytoplankton in nutrient-poor environments [11, 12]. In contrast, larger-sized phytoplankton can sustain higher rates of biomass-specific production rate in nutrient-rich waters and less tightly controlled by grazers [13–15]. These constraints lead to different temporal and spatial distribution patterns of smaller and larger phytoplankton. In general, nano- and microphytoplanton (> 2 μm) dominate in productive areas, whereas the pico-phytoplankton (0.2 to 2 μm) dominate the autotrophic biomass and the production of oligotrophic regions [2, 11, 16]. Several studies recently reported that temperature directly affects the size structure of phytoplankton assemblages, such that picophytoplankton contributes more phytoplankton biomass in warm water [16, 17]. Therefore, how the temperature and other factors influence different size group community assembly is an exciting but underexplored topic.

Eukaryotic plankton community assembly is under the influence of many complex factors e.g. selection, mutation, disperse, and gene flow [18]. An increasing number of studies have focused on the importance of two contrasting processes in determining the structure of the eukaryotic plankton community. One is the deterministic process (environmental factors and species interactions-related) [18], and the other is the stochastic process (dispersal, immigration-related) [19, 20]. Most recent studies reported a significant relationship between microeukaryotic community composition and deterministic processes in various aquatic ecosystems. For example, Sunagawa et al. (2015) showed that the sunlit plankton community’s vertical structure was mostly influenced by temperature rather than geography at a global scale [4]. Another study reported that the environmental variable could explain 34.76% of the entire microeukaryotic community variation, whereas spatial factor only explained 18.62% of the variation in the northwestern Pacific Ocean [21]. For benthic microeukaryotes in marine sandy beaches, a recent study found that abundant and rare microeukaryote subcommunities exhibited a stronger response to environmental factors than spatial factors [7]. However, a study in the East China Sea and the South China Sea indicated that abundant subcommunities were strongly influenced by dispersal limitation [6]. It remains unclear whether the microeukaryotic compositions including larger-sized (nano- and micro-plankton) and small-sized (picoplankton) biospheres are influenced more by the deterministic or the stochastic process in the open ocean.

In this study, we used high-throughput DNA metabarcoding to compare the species composition and diversity of the eukaryotic plankton community at the continental shelf (C6 station) and continental slope (C9 station) in the northern South China Sea. The first objective was to characterize the eukaryotic community structure in different habitats and interspecies network among eukaryotic plankton. Then with the data, we aimed to examine two hypotheses regarding eukaryotic plankton community assembly: 1) the deterministic process and the stochastic process may have differential importance in structuring larger-sized and smaller-sized subcommunities in the eukaryotic plankton community, and 2) the smaller organisms can disperse more broadly than larger organisms (the size-dispersal hypothesis). Our results provide novel insights into the influencing factors and mechanisms in the assembly of large-sized and small-sized biospheres in the microeukaryotic community in the northern South China Sea, a subtropical oceanic province.

## Results

### Diversity and distribution of eukaryotic plankton

We recovered 2,148,738 high-quality sequences from the 49 samples collected from C6 and C9 stations (Figure 1), which were clustered into 2,463 OTUs at 97% sequence similarity level (Table 1). The rarefaction curves were roughly saturated for most samples (Figure S1), indicating that most of the microeukaryotic plankton taxa had been retrieved from the two stations. The numbers of OTUs varied from 370 to 1044 per sample (Table S1). Alpha diversity Shannon index for each sample ranged from 2.38 to 4.88 at station C6 and from 2.99 to 5.10 at C9 (Table S2, Figure S2), indicating a shelf to slope increasing trend. The metacommunity was overwhelmingly dominated by alveolates and opisthokonts in terms of taxon number and abundance (Figure 2). Among the total of 2,463 OTUs, 1,461 belonged to Alveolata (relative abundance = 44.76%), 302 Opisthokonta (31.08%), 279 Rhizaria (13.79%), 240 Stramenopiles (2.92%), 53 Incertae_Sedis (2.81%), 3 Amoebozoa (0.02%), 40 Archaeplastida (0.56%), and 3 Excavata (0.01%) (Figure 2). In the entire dataset, 2,085 OTUs, with 1,326,555 sequence reads (62.74%), dominated the 3-200 μM size fraction, whereas 2,193 OTUs with 822,183 sequence reads (38.26%) dominated the 0.22-3 μM size fraction (Table 1).

**Fig 1.**
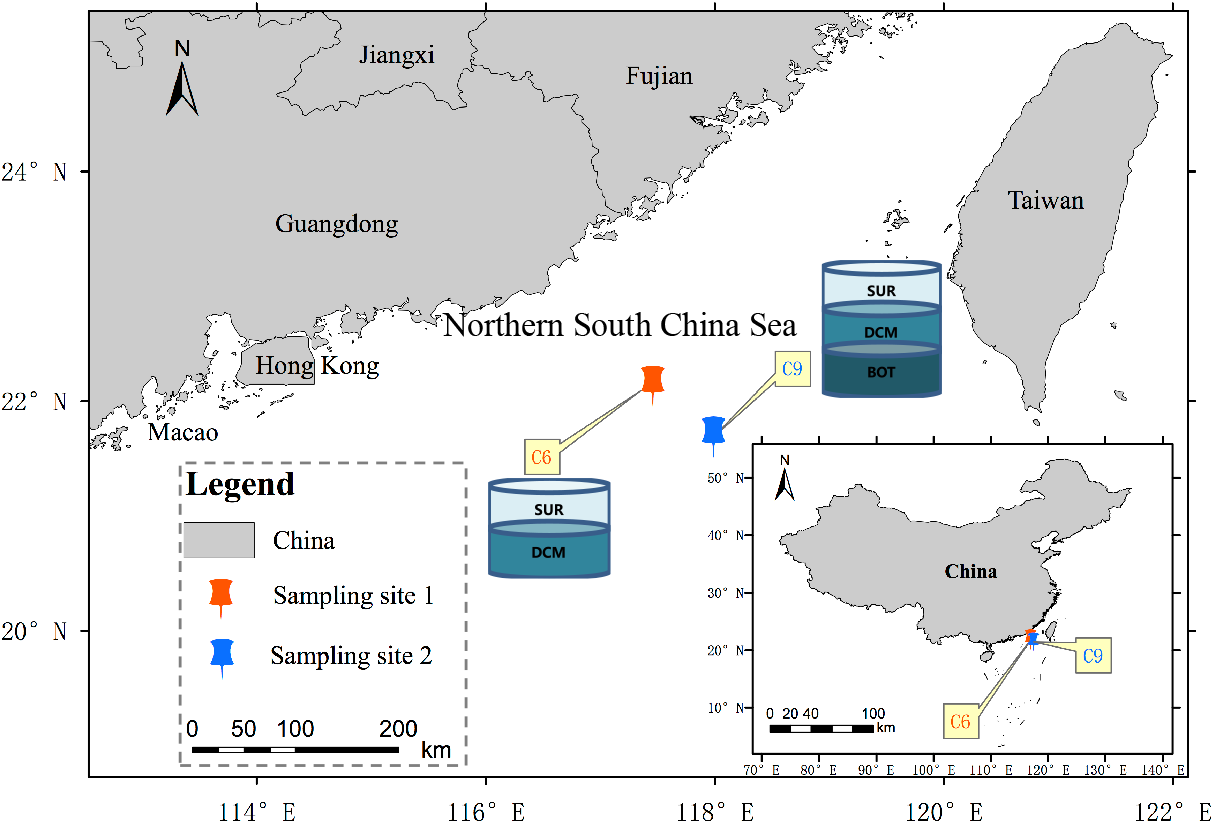
Locations of the sampling stations. Station C6 (117.46°E, 22.13°N) is on the continental shelf, with a depth of 77 m. Station C9 (117.99°E, 21.69°N) is on the continental slope, with a depth of 1369 m. SUR: surface (0-3 m); DCM: deep chlorophyll maximum (35 m at C6 and 75 m at C9); BOT: depth named bottom (150 m).

**Table 1.**
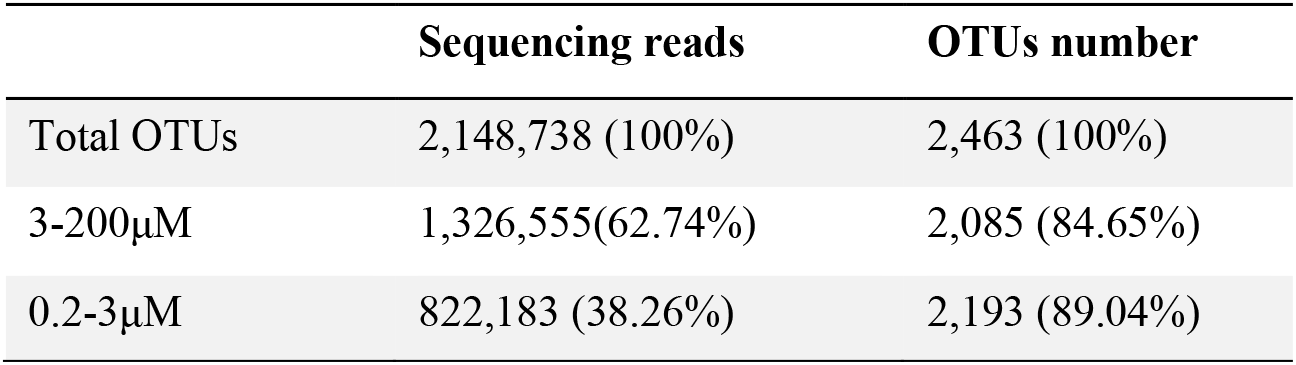
Statistics of total, 3-200μM, and 0.2-3μM OTUs of the metacommunity.

|  | Sequencing reads | OTUs number |
| --- | --- | --- |
| Total OTUs | 2,148,738 (100%) | 2,463 (100%) |
| 3-200µM | 1,326,555(62.74%) | 2,085 (84.65%) |
| 0.2-3µM | 822,183 (38.26%) | 2,193 (89.04%) |

**Fig 2.**
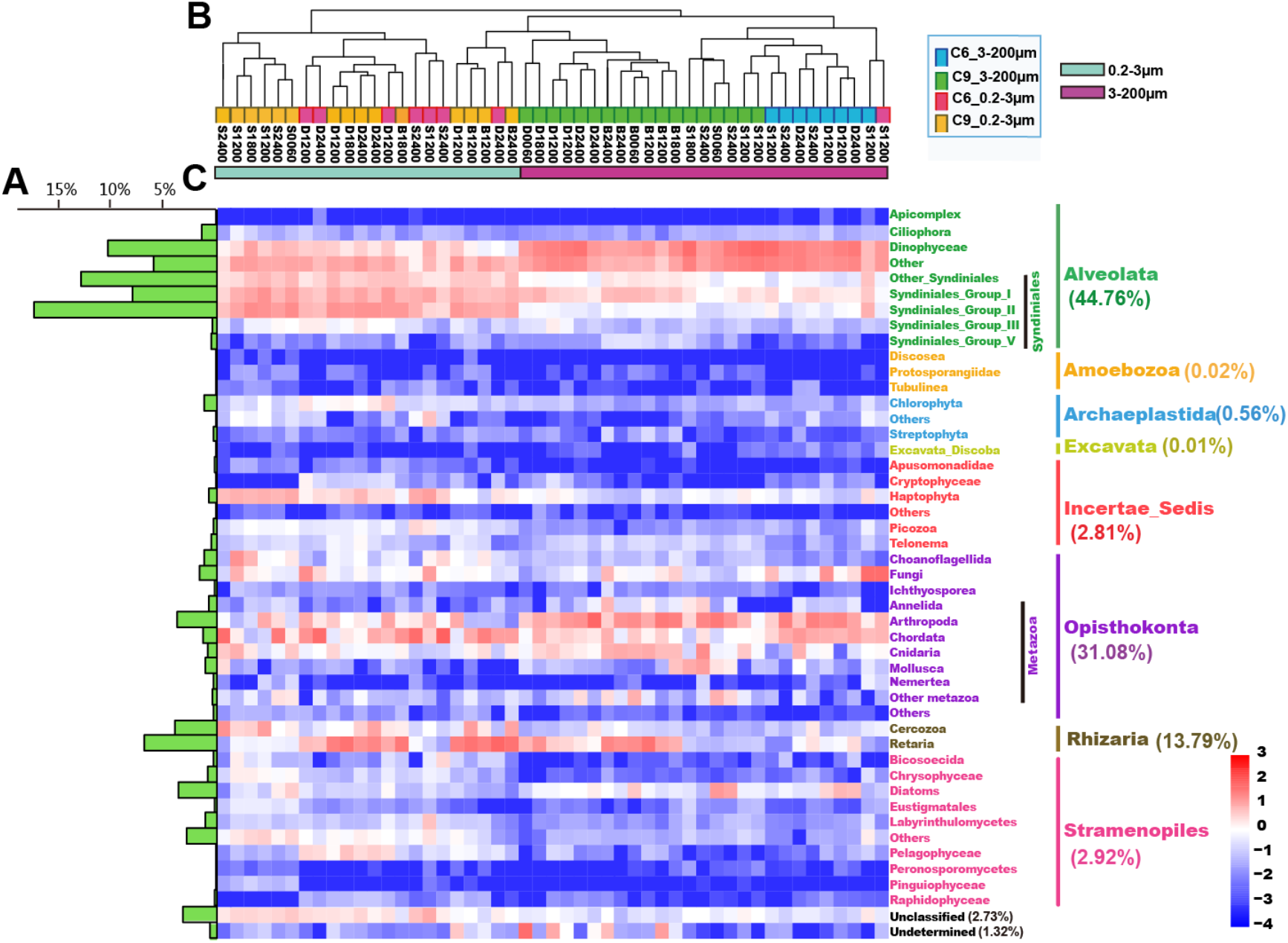
Phylogenetic distribution and relative abundance of the microeukaryotes. A) The relative abundance of sequencing reads in the 47 lineages detected within the entire dataset. B) Dendrogram cluster for 49 samples based on Bray-Curtis similarity. The diversity color coding at the branch end based on their sampling size fractions and stations. C) Heatmap of the relative abundance of the 47 lineages within each sample.

The eukaryote communities showed a distinct size-dependent distribution pattern between the two sampling stations (Figure 2). The two size fractions formed separate clusters, and in the larger-size cluster, samples were further grouped by station, whereas in the other cluster, samples from different stations mingled. This pattern indicated that the picoplankton assemblage was more similar between the two stations than the nano-micro plankton assemblage. The smaller-size fraction contained a greater proportion of shared OTUs across stations and depths than did the larger size fraction (P < 0.01), indicating that both horizontal and vertical dispersal abilities were higher for the smaller-size fraction than the larger-size fraction (Figure 3). Furthermore, PCA analysis revealed that the bulk OTUs were distinctly separated by size fraction (22.46%), and the differences were confirmed by ANOSIM statistics analysis (R = 0.77, P < 0.01) (Figure 4A). Moreover, the Venn diagram showed the 881 OTUs (36%) were present in all the three depths we sampled, including 745 OTUs (36%) in 3-200 μM and 748 OTUs (34%) in 0.22-3 μM (Figure 4B).

**Fig 3.**
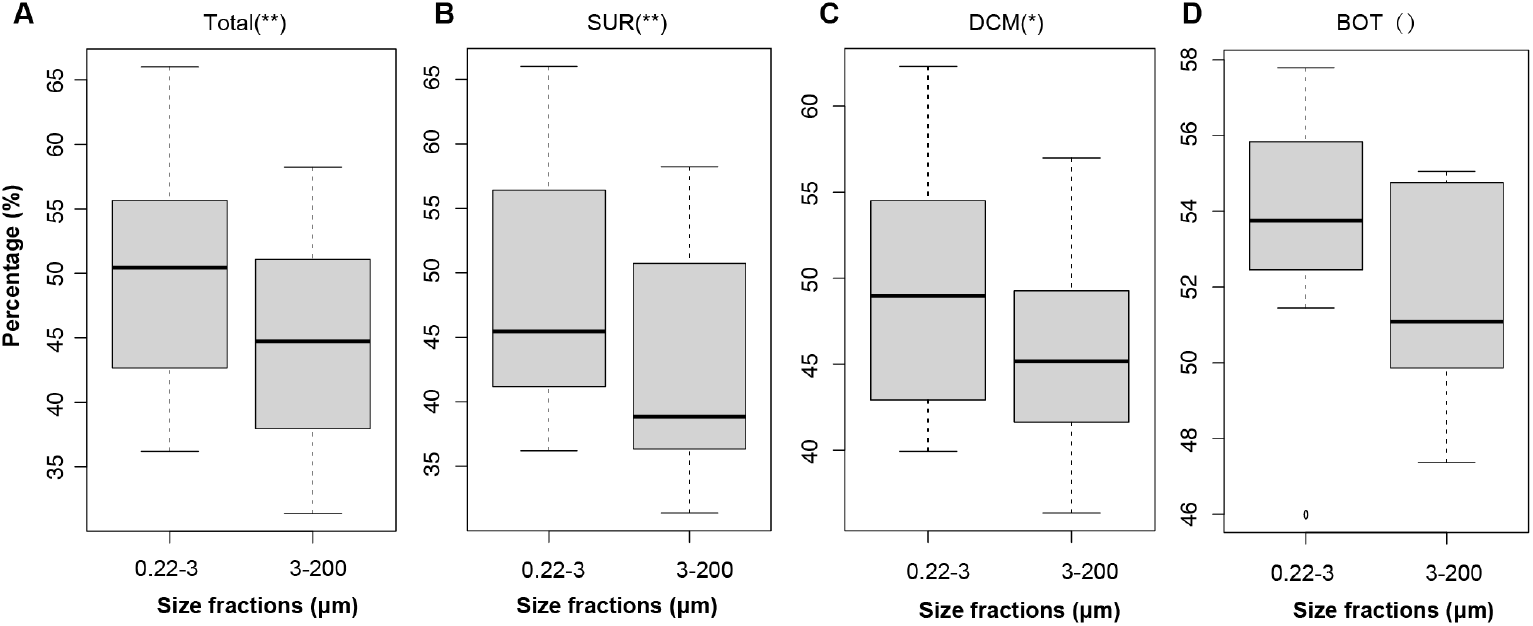
Boxplots showing mean shared proportions of OTUs of eukaryotic community in three sampling depths. A greater proportion represents a greater average dispersal ability of organisms in the group. **A**) The bulk shared proportions for two size organisms including three sampling depths. **B**, **C**, **D**) Comparison of two size organisms in each water depth. Significant differences were marked with an asterisk in brackets (**: P < 0.01; *: P < 0.05).

**Fig 4.**
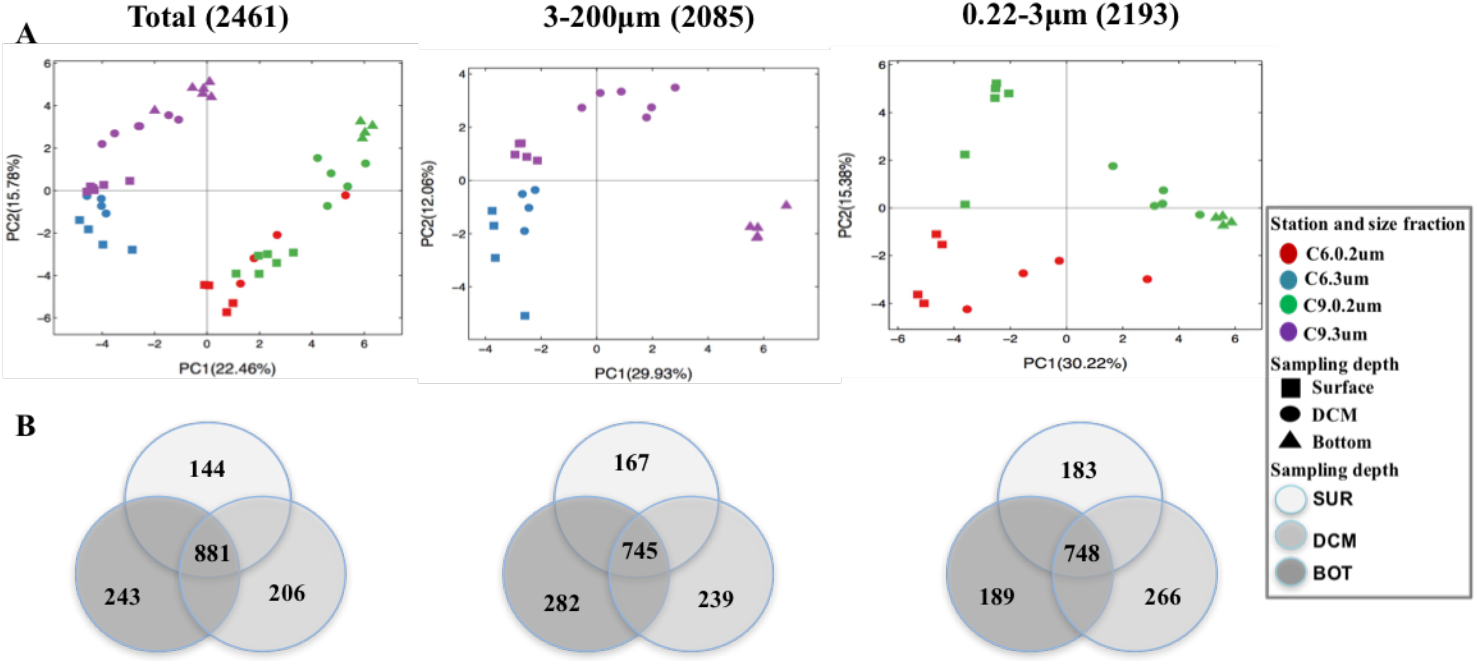
Comparison of microeukaryote communities from continental shelf (C6) and slope (C9) stations. A) Grouping of local communities according to compositional similarity using Principal Component Analysis (PCA). Each symbol represents one sample or microeukaryotic community. Samples in the two stations with two size fractions and three depths were marked in different colors and/or shapes. B) Venn diagrams showing the number of OTUs that are unique and shared between three depths.

### Differences in community structure between C6 and C9 stations and different depths

We obtained 1,746 OTUs in the C6 station, 370 to 1,011 per sample, and 2,300 OTUs in the C9 station, 476 to 1,044 per sample (Table S1). Analysis of similarity (ANOSIM) showed that the communities of C6 and C9 stations were significantly separated (R = 0.220, P < 0.01) (Table 2). Group comparison based on sampling depths and size fractions showed significant difference sampling depths (P < 0.05, Table 2). Moreover, diversity indices were higher at C9 station than C6, significantly so for indices of Sobs, Chao, Ace, and Shannon for surface samples (Mann-Whitney *U* test, P < 0.05) and for Shannon index for DCM sample (Mann-Whitney *U* test, P < 0.05). These indices consistently indicated that station C9 was more biodiverse than C6 (Table S2, Table S3). Lineage wise, the relative abundances of Syndiniales (parasitic dinoflagellates), marine Ochrophytes (MOCH), Haptophyta, and Choanoflagellida were higher at C9, whereas that of Metazoa, especially in smaller size fraction, was lower at C9 (Mann-Whitney *U* test, P < 0.05; Figure S3).

**Table 2.**
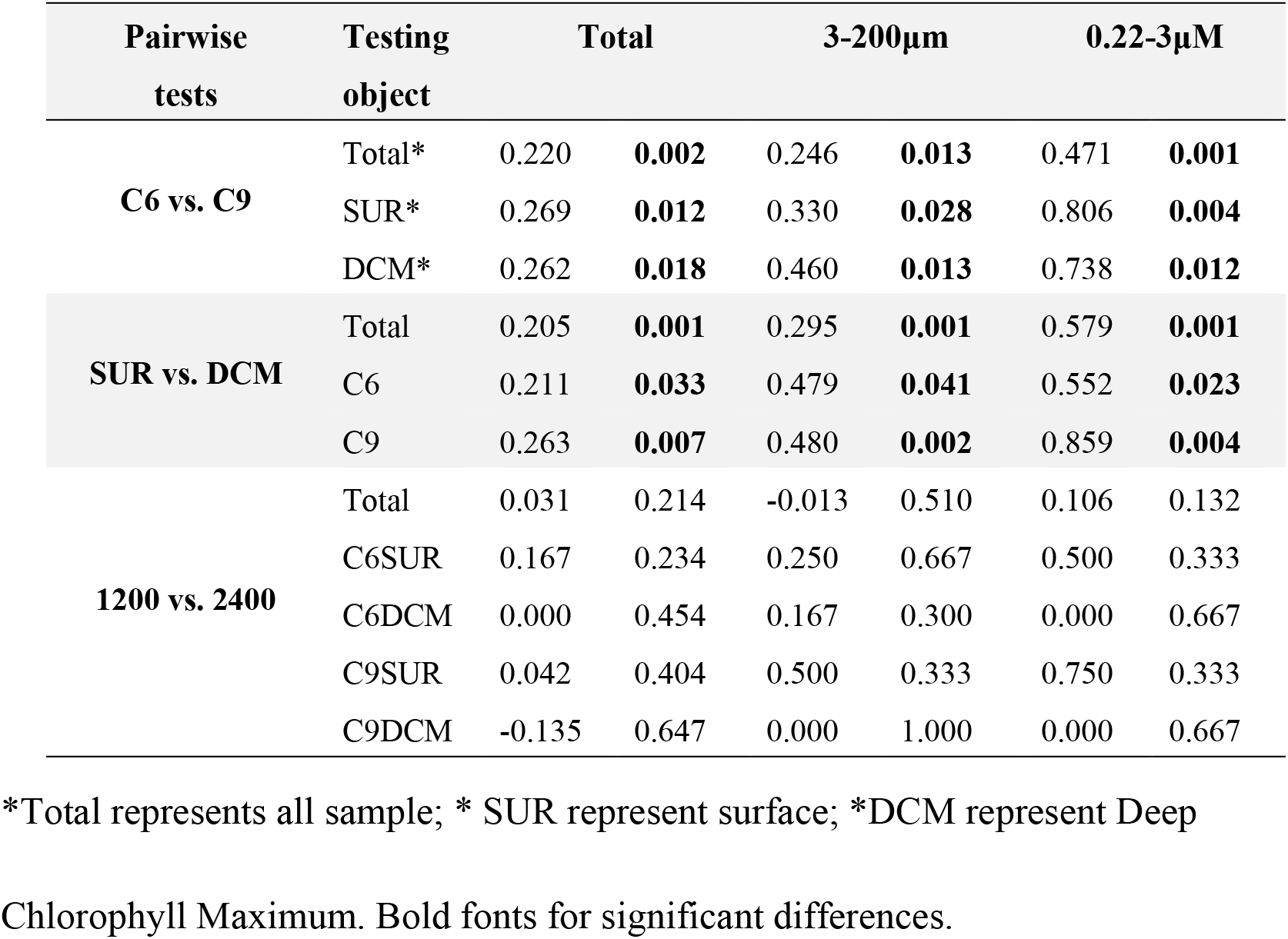
Analysis of similarities (ANOSIM) of total community, larger size (3-200μM) and smaller size (0.22-3μm) sub-communities among different regions, depths and sampling times.

| Pairwise tests | Testing object | Total |  | 3-200µm |  | 0.22-3µM |  |
| --- | --- | --- | --- | --- | --- | --- | --- |
| C6 vs. C9 | Total* | 0.220 | **0.002** | 0.246 | **0.013** | 0.471 | **0.001** |
|  | SUR* | 0.269 | **0.012** | 0.330 | **0.028** | 0.806 | **0.004** |
|  | DCM* | 0.262 | **0.018** | 0.460 | **0.013** | 0.738 | **0.012** |
| SUR vs. DCM | Total | 0.205 | **0.001** | 0.295 | **0.001** | 0.579 | **0.001** |
|  | C6 | 0.211 | **0.033** | 0.479 | **0.041** | 0.552 | **0.023** |
|  | C9 | 0.263 | **0.007** | 0.480 | **0.002** | 0.859 | **0.004** |
| 1200 vs. 2400 | Total | 0.031 | 0.214 | -0.013 | 0.510 | 0.106 | 0.132 |
|  | C6SUR | 0.167 | 0.234 | 0.250 | 0.667 | 0.500 | 0.333 |
|  | C6DCM | 0.000 | 0.454 | 0.167 | 0.300 | 0.000 | 0.667 |
|  | C9SUR | 0.042 | 0.404 | 0.500 | 0.333 | 0.750 | 0.333 |
|  | C9DCM | -0.135 | 0.647 | 0.000 | 1.000 | 0.000 | 0.667 |

\*Total represents all sample; \* SUR represent surface; \*DCM represent Deep
Chlorophyll Maximum. Bold fonts for significant differences.

Comparing the diversity across the three sampling depths, we found that the deepest depth (BOT) harbored the highest diversity, whereas the surface had the lowest (Table S4), for all the four diversity indices (Sobs, Chao, Ace, and Shannon) (Kruskal-Wallis *H* test, P < 0.05). Notably, the DCM layer in C9 station had a higher diversity than BOT layer in the smaller size fraction (Table S4). This indicated that depth was an influencing factor for the eukaryotic community (Table S5). In support of this, ANOSIM analysis showed significant differences between the SUR and DCM (R = 0.205, P < 0.05). In addition, we found that Pelagophyceae, Polycystinea, Cryptophyceae, RAD, and Chlorophyta were more abundant in DCM, whereas Dictyochophyceae, Chrysophyceae, and Haptophyta were more abundant in SUR (Mann-Whitney *U* test, P < 0.05, Figure S4), indicating more pronounced depth-wise variation in these eight lineages than others. Comparing between-station differences with between-depth differences, community dissimilarities were 0.205 between SUR and DCM (P < 0.05) and 0.220 (P < 0.05) between C6 and C9, indicating a greater difference between the two sampling sites (Table 2). The opposite was found for larger-size and smaller-size subcommunities, in which a greater difference occurred between the two sample depths (Table 2).

### Network analysis of microeukaryotes and environmental factors

Regarding the interactions between microeukaryotes, more significant correlations were found in the smaller-size fraction than the larger-size fraction (Figure 5). In the 0.22-3 μm fraction, RAD and Polycystinea showed a negative correlation with most eukaryotes (P < 0.05), whereas most other groups, including Syndiniales, Haptophyta, Dictyochophyceae, MAST, MOCH, and Chrysophyceae, showed significant positive correlations with other eukaryotes (P < 0.05; Figure 5). In the 3-200 μm fraction, Metazoa were negatively correlated with Dinophyceae and Haptophyta (P < 0.05), whereas Dinophyceae, Haptophyta, and MOCH were positively correlated with most other eukaryotes (P < 0.05). Regarding environmental factors, we found no significant correlation between NO_2_^-^ concentration and any groups of organisms, whereas NO_3_^−^+NO_2_^−^ concentration showed a significant correlation with most lineages, e.g. Dinophyceae and Haptophyta. In addition, depth, temperature, and salinity showed more correlations with eukaryotes than other nutrient factors (Figure 5).

**Fig 5.**
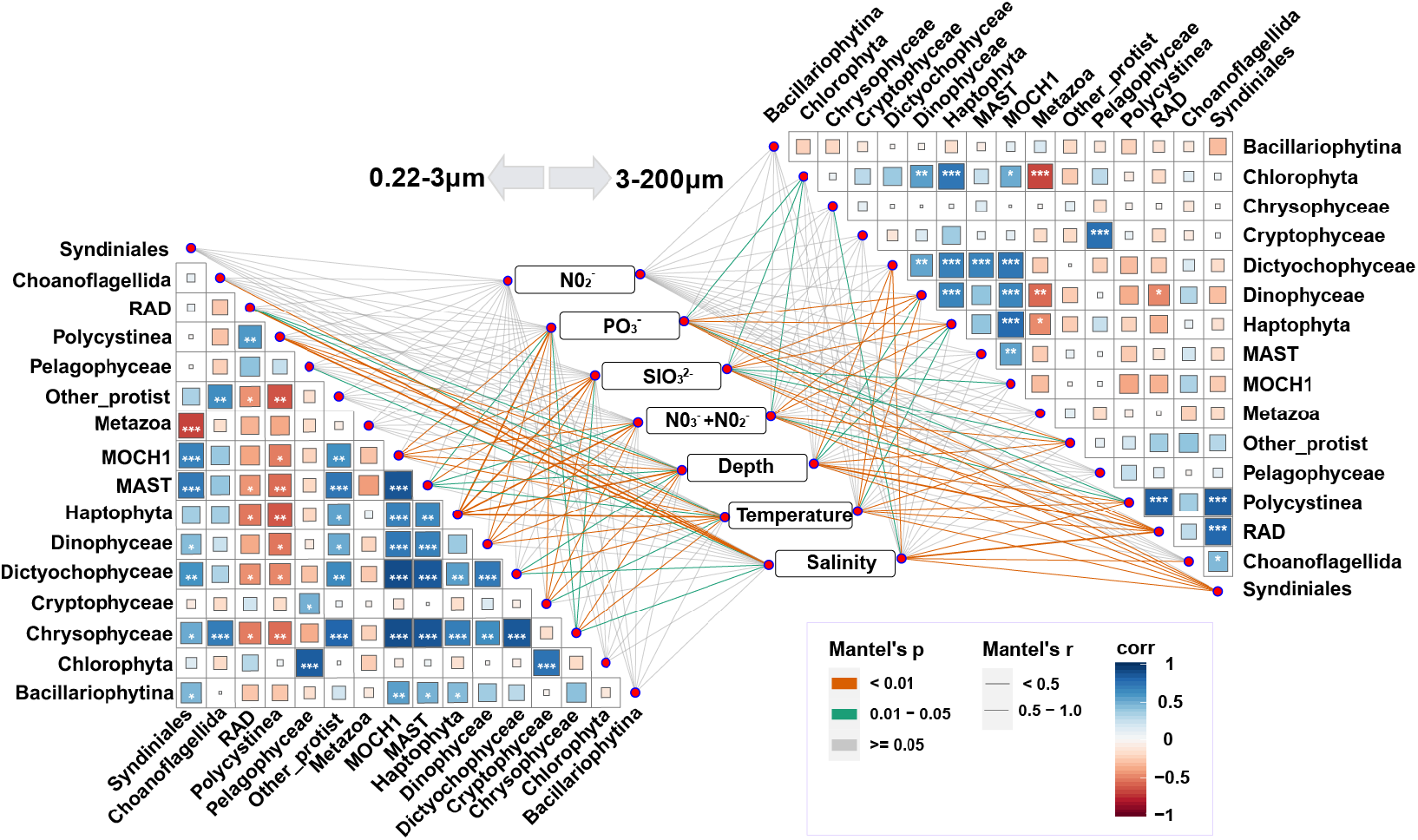
Interactions across microeukaryotes and their relationships with environmental factors. The thickness of the line represents the correlation between environmental factors and taxon abundance, and the color of the lines represents the significance level of the correlation (see color scale and thickness scale on the upper and middle right). Fill color strength of the squares represents the correlation of selected major taxa, from orange (negative interaction), white, to blue (positive interaction) (see color scale on the lower right). Asterisks indicate significance (*: P < 0.05; **: 0.05 < P < 0.01; ***: P < 0.01).

### Diel variation pattern

Two or four diel samples were collected at stations C6 and C9, respectively. Analysis results indicated that 10%-36% (average = 19%) of all OTUs exhibited two diel dynamics patterns (Figure S5). One was characterized by low abundance at noon and high abundance at midnight (pattern 1), and the other just the opposite (pattern 2). A total of 610 OTUs belonged to pattern 1 and 1033 OTUs pattern 2, indicating a substantial the diel (likely circadian) variation influencing plankton community structure. Furthermore, depth, temperature and salinity showed a significant correlation with pattern 1 and pattern 2 (Figure S6). However, although the diel dynamics manifested in 19% of all OTUs, ANOSIM analysis showed that the entire community displayed no significant difference between 1200 and 2400 (P > 0.05, Table 2).

Metazoa and Dinophyceae, which dominated the 3-200 μm fraction, changed with time in relative abundance, especially in the surface layer. In the 0.22-3 μm fraction, the relative abundances of the dominant groups Metazoa, Syndiniales, Haptophyta, and Polycystinea also varied temporally but with no consistent patterns (Figure 6). Between time points 1200 and 2400, Metazoa, MAST, and Unassigned group showed significant differences (Mann-Whitney *U* test, P < 0.05, Figure S7).

**Fig 6.**
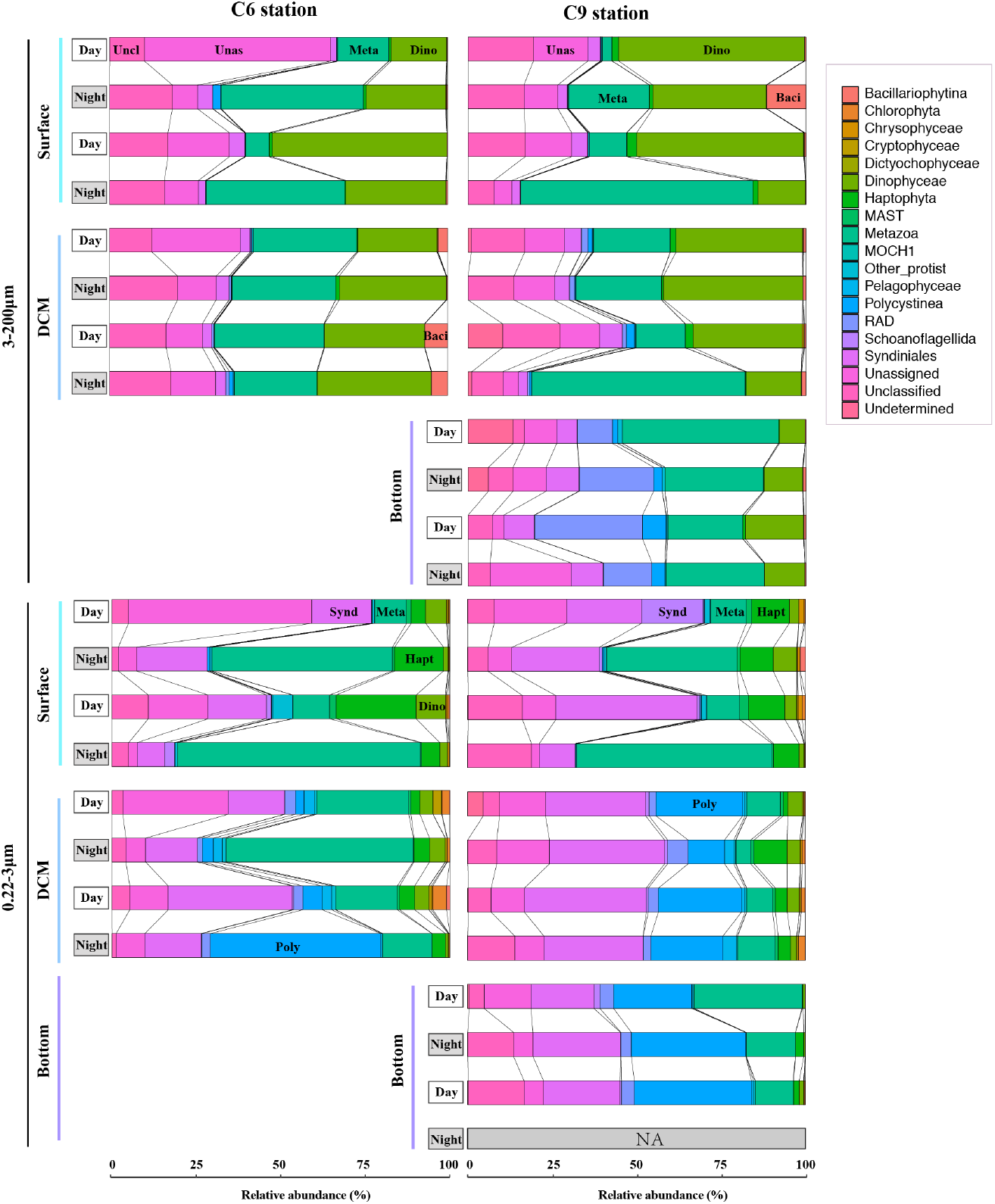
Relative abundance and diel variation of major eukaryotic taxa. Relative abundance data were from four successive time points across different stations, different depths and different size fractions.

### Correlations of eukaryotic community with environmental factors and geographic locality

Results from VPA showed that spatial and environmental factors together explained 25%, 39% and 47% variations of the entire community, larger-size community, and smaller-size community, respectively (Figure 7). RDA and forward selection showed that temperature and salinity were the main explanatory variables among environmental factors, which accounted for 19% variation for the entire community, 28% variation for the larger-sized community, and 32% variation for the smaller-sized community. Further analysis revealed that temperature explained more pure variation than did salinity in the bulk community (5% vs. 4%), larger-size community (8% vs. 5%), and smaller-size community (9% vs. 7%). Mantel and partial Mantel tests further confirmed that the bulk community, larger-size community, and smaller-size community were primarily governed by environmental factors rather than spatial factors (Table 3).

**Fig 7.**
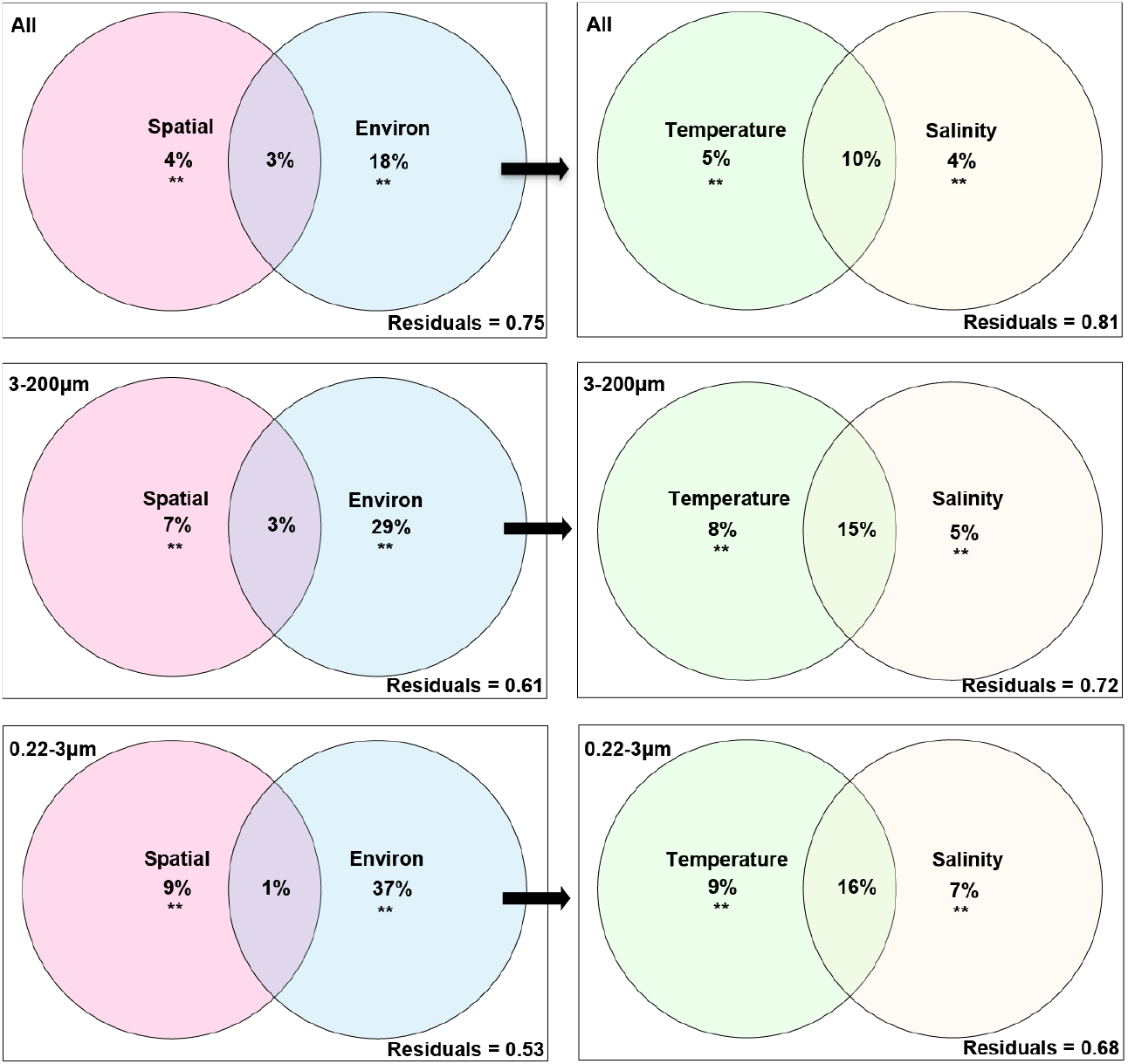
Venn diagrams of variation partitioning analysis showing the effects of spatial and environmental factors on the microeukaryotic communities. The percentage of variation explained by each factor, including unique, shared and unexplained (Residuals) is shown in corresponding positions in diagram. ANOVA permutation tests were calculated on the pure variation. ^**^P < 0.01.

**Table 3.**
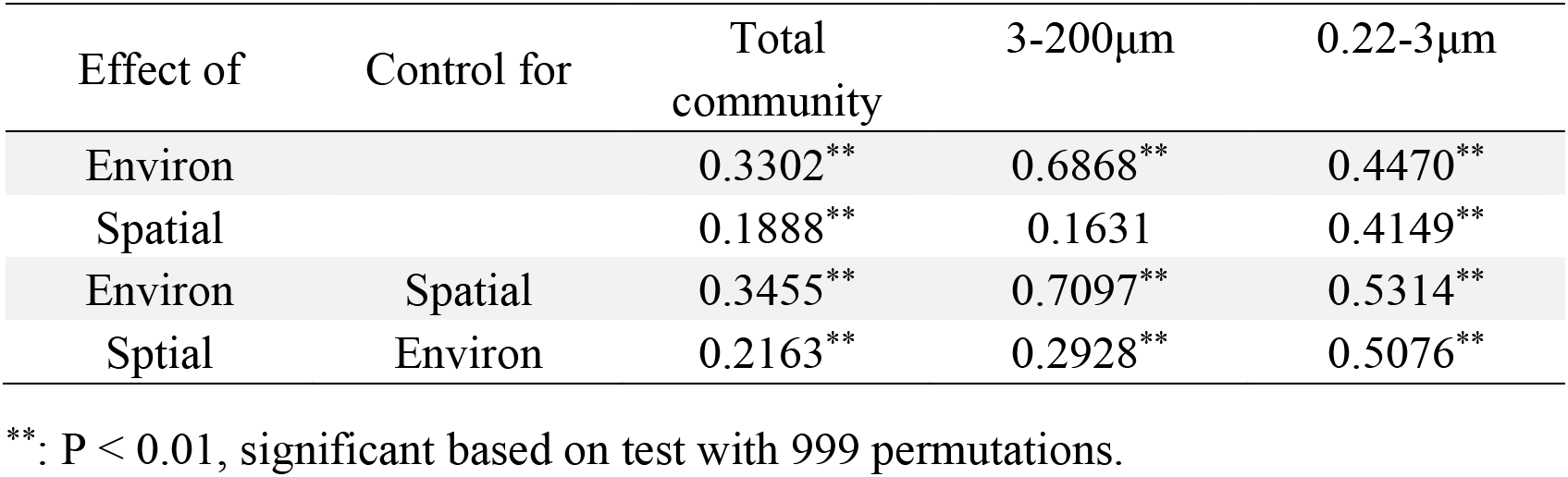
Mantel and partial Mantel tests for the correlation between community similarity and spatial and environmental variables using Pearson’s coefficient.

### Neutral assembly of eukaryotic community assemblages

The neutral model explained a small fraction of the variation (19%) in the occurrence frequency of different OTUs in the bulk eukaryotic community (R^2^ = 0.19, Figure 8A). Comparatively, the model explained a much higher fraction of variation for the smaller-sized community (R^2^ = 0.242) than the larger-sized community (R^2^ = 0.159) (Figure 8B, C). The model also estimated a higher migration rate in the smaller-sized community (m = 0.076) than in the larger-sized community (m = 0.073). Furthermore, a higher percentage of OTUs in the smaller-sized community (73.4%) fell within the neutral distribution range than did the larger-sized community (70.5%) (Figure 8B, C), indicating that the smaller-sized plankton subcommunity assembly fit the neutral model better than the larger-sized subcommunity.

**Fig 8.**
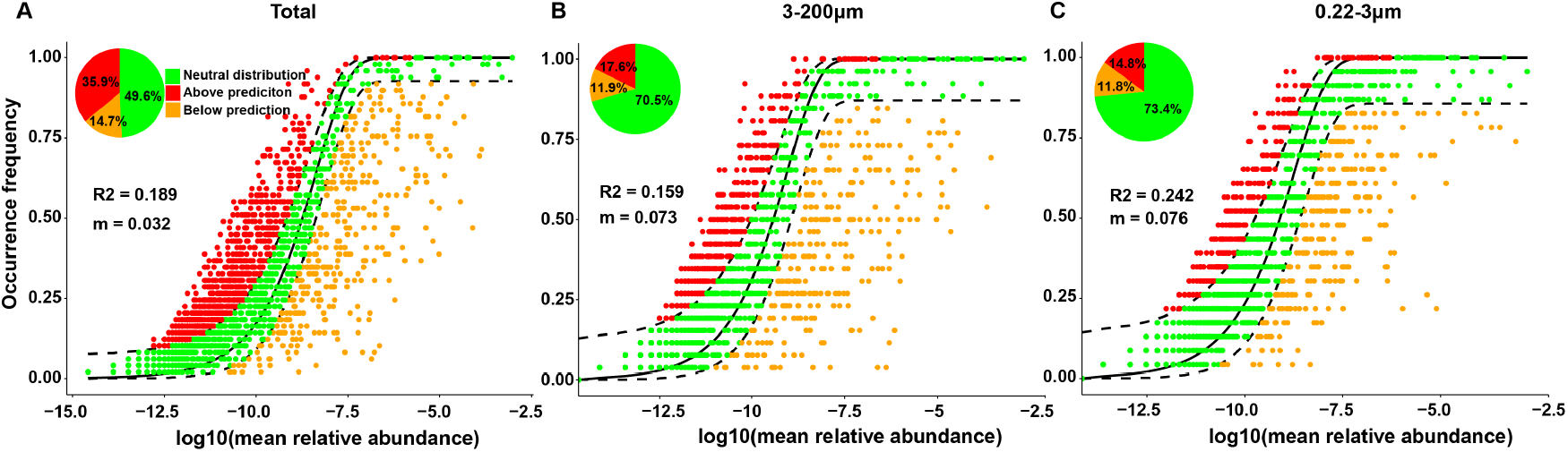
Fit of the neutral model on the assembly of eukaryotic community (A) including larger and small size communities (B, C). R2: represent the goodness-of-fit of neutral model; m: represent estimated migration rate. The sold black lines represent the best fit to the neutral model, and the dashed black lines represent 95% confidence internals around the model prediction. The pie charts depict the richness of the neutral distribution, above prediction and below prediction.

## Discussion

### Eukaryotic diversity and distribution in continental shelf and slope

In recent years, the assembly of microeukaryotic communities in global oceans has gained an increasingly deep understanding across increasingly broader geographic scales [4, 5]. On a global scale across a latitudinal gradient, temperature was found to be the primary environmental factor shaping plankton’s diversity [22]. At smaller scales, how the community assembly differs between different oceanic provinces (e.g. continental shelf and slope) has been understudied and less well understood. Our study addresses the gap of effort and provides insights into the processes and factors shaping the communities along a shelf-slope gradient in a subtropical seascape. Our effort documents all eukaryotic super-groups that have been documented and reveals variations and dynamics of the plankton community in different habitats (Figure 2).

Our results showed that Alveolata (mainly dinoflagellates, including the parasitic order Syndiniales) and Opisthokonta (mainly Metazoa) were dominant both in diversity and abundance (Figure 2). The same has previously been observed from other ecosystems, where the euphotic zone was sampled [5, 21]. Our multi-depth sampling scheme further indicated that alveolates and opisthokonts were dominant in DCM and BOT layers too, where light intensities were estimated to be 0.75 μE m^−2^ s^−1^ and 0.02 μE m^−2^ s^−1^ in C9 station based on a remote sensing model [23]. Syndiniales is a highly diverse lineage, containing parasites of fish eggs (group I) and parasites of dinoflagellates (group II, mainly the genus of Amobephrya) [24]. They are an abundant constituent of eukaryotic plankton, particularly picoplankton, in various ecosystems [24–27]. For other dinoflagellates (Dinophyceae), their high relative abundances detected in 18S rDNA libraries may be in part due to their generally immense genomes (~1-250 Gbp) and high 18S rDNA copy numbers [28, 29], in contrast to most other algae whose genome sizes typically range from tens to hundreds of million bp (Mb) [30]. However, dinoflagellates are an important group of protists with a wide distribution and strong adaptive ability [31], and their dominant abundance in oceanic plankton has also been observed using microscopic analyses [32, 33]. The dominance of Syndiniales is even less attributable to genome size because the parasitic lineage generally has a small genome size (e.g. *Amoebophyra ceratii* genome size is about 100 Mbp; [34]).

Metazoan’s prominent contribution to plankton is notable. As many of the organisms are larger than 200 μm in body size, the result suggests a high abundance of eggs, spore or larvae of large-sized zooplankton and other animals or small-sized metazoan (e.g. Orthonectida), which can pass through 200μm bolting cloth and contribute to the plankton assemblage [35]. The top three metazoans in our samples were Copepoda, Collodaria and Cnidaria. In previous metabarcoding analyses of marine plankton from other ecosystems, these three lineages were also found to be the most abundant metazoan [5].

In the present study, more OTUs were found in the continental slope (2,300) than continental shelf (1,746), consistent with the current notion that the offshore biodiversity is higher than nearshore counterparts [36]. Previous studies showed that haptophytes appeared to be more dominant in the open ocean than coastal areas [37, 38]. In the present study, the abundance of the haptophytes and Syndiniales, MOCH, and Choanoflagellida were higher on continental slope than on continental shelf. Furthermore, we found that the biodiversity increased as the water depth increase, with the highest diversity retrieved from BOT (Table S2, Table S4).

### Factors controlling the assembly of the eukaryotic plankton community

Both the spatial and environmental processes play significant roles in shaping the distributions of eukaryotic plankton communities in the ocean [21, 39]. Unlike the extensive effort on marine microbial (prokaryotic) communities, studies on the effect of spatial and environmental factors on eukaryotic plankton community assembly are still relatively rare. Our study evaluated potential controlling factors and the relative contributions of spatial and environmental factors in shaping the eukaryotic plankton community. We found that both environmental and spatial processes played a significant role in structuring eukaryotic plankton assemblages, including larger-sized and smaller-sized communities, from the continental slope to the shelf (Table 3, Figure 7). Furthermore, we found that the environmental factor (mainly temperature and salinity) had a greater impact than the spatial factor, indicative that the deterministic process is more important in shaping the community than the stochastic process in our study oceanic area. This may be due to the relatively short distance between the two sampling stations (73.3 Km) and between depths, although station appeared to have a greater influence on community structure than depth (surface and DCM) for the bulk community, consistent with the greater inter-station distance than inter-depth distance (Table 2). Alternatively, the active surface currents forced by the East Asia monsoonal winds in this seascape may weaken dispersal limitation [40].

For oceanic waters, diel variation has been documented widely (e.g. *Prochlorococcus*, *Synechococcus*, *Resultor micron*, *Heterosigma akashiwo*) [41–44]. Consistent with these findings, we discovered that on average 19% of the detected OTUs in each divided group, including metazoa and MAST groups, showed clear diel patterns in relative abundance (Figure S7, Figure S9). These patterns can stem from diel rhythms of cell division, grazing activity, and diural vertical migration [45–48]. However, no significant diel pattern was observed for the entire microeukaryotic community, larger-sized and smaller-sized communities (Table 2). This was because the combination of opposite diel patterns of different lineages led to the diel variation’s cancellation. Therefore, bulk community-level analysis can miss the diel variation occurring in a substantial component of the community.

For environmental factors, it has been reported that temperature is a more important factor shaping microeukaryotic community composition and diversity than other factors on the global scale [4, 21, 22], and a greater species diversity occurs in the low-latitude ocean because of high temperatures [49]. In addition, a large-scale meta-analysis indicated that salinity was one of the major determinants across different habitats and significantly correlated with microeukaryotic community [21, 50]. Consistent with these previous results, we found that temperature and salinity explained more variation of eukaryotic plankton community assembly than other environmental factors in our study area (Figure 7). Although other environmental factors explained smaller community variation, some of them, e.g. nitrogen and phosphorus, exhibited a strong correlation with some taxonomic groups, indicating that the nutrient effects are more lineage-specific (Figure 5). Notably, only temperature differed significantly between the shelf and the slope stations (Figure S9).

### Dominant role of stochastic process in microeukaryotic community assembly

Several studies reported that a null or neutral-based model-based analysis could enhance our ability to understand the relative importance of different ecological processes underlying microbial community assembly [9, 51, 52]. As a neutral-based process model, the neutral community model is useful for inferring stochastic processes acting on community assembly [53]. Our results showed that the stochastic processes explained 18.9% variation of the bulk microeukaryotic community assembly, 15.9% variation for larger-sized community, and 24.2% variation for the smaller-sized community (Figure 8). Different sized eukaryotic plankton may have fundamentally different characteristics and ecological roles [11]. In the present study, we found more OTUs but less abundance in the smaller-size community than the larger-sized community (Table 1). Furthermore, the value of neutral model parameter *R^2^* and m were slightly higher in the smaller-size community than the larger-sized community, indicating that dispersal between the shelf and slope study sites was higher for the picoeukaryotic plankton community than the nano-micro eukaryotic plankton community. These results indicate that the stochastic process’s influence, at least in terms of dispersal, seems to be stronger for the picoeukaryotic than the nano-micro eukaryotic plankton community.

In addition, we found the larger-sized plankton assemblages were less similar between stations and depths than smaller-sized plankton assemblages (Figure 2B), suggesting a stronger dispersal or niche limit for the larger-sized community. Size-dispersal and size-plasticity hypotheses have been proposed, which postulate that larger organisms are more dispersal- or niche-limited than smaller species [54–56]. Consistent with the neutral model result and size-dispersal hypothesis, a greater average dispersal ability was found in the smaller-sized plankton community than larger-sized plankton (Figure 3). To our knowledge, this is the first documentation of evidence supporting the size-dispersal hypothesis for eukaryotic plankton. In contrast, the niche breaths were estimated to be the same between the two size fractions (Figure S8), indicating that the size-plasticity hypothesis was not suited to explain the differential distribution patterns of picoeukaryotic plankton and nano-micro eukaryotic plankton assemblages in our study.

### Effects of interspecies interactions on the dynamics of the eukaryotic plankton community

Although our analysis revealed that deterministic and stochastic processes explained some of the community variations, some variation remained unexplained, which might be attributable to unmeasured biotic and abiotic factors. Previous research showed that drift and mutation also created and maintained microbial biogeographic patterns on inseparable ecological and evolutionary scales [39]. Furthermore, studies found that the interactions among microbes, including predation, parasitism, mutualism, competition, and cross-feeding, are also responsible for shaping community structure [57, 58]. In the present study, we found that RAD and Polycystinea were negatively correlated with other eukaryotic plankton, indicating a potential competitive relationship between these lineages. Syndiniales were negatively correlated with metazoa, suggesting a possibility that Syndiniales have a parasitic relationship with metazoa. This is not a surprise because this diverse and abundant marine protist lineage is exclusively parasitic [24–27].

### Concluding Remarks

The present study documented higher biodiversity of eukaryotic plankton in the continental slope than the continental shelf and increased biodiversity with depth in the subtropic northern South China Sea. We also found that the deterministic process was more important than the stochastic process in the assembly of the eukaryotic plankton communities at all levels, including the bulk plankton community, the picoeukaryotic plankton subcommunity, and the nano-micro eukaryotic plankton subcommunity. However, a substantial part of the community variability remains unexplained, indicating that other unmeasured factors also contribute to the community’s assembly in these two environments. The entire community was structured by multiple factors with importance degree: environmental factors (temperature + salinity) > spatial factors > water layers > sampling times. Among environmental factors, temperature was the main influencing factor in shaping the eukaryotic plankton community, suggesting that nutrients in general were not as important as previously thought in shaping the eukaryotic plankton community in this oceanic province. Furthermore, the smaller-sized fraction of the eukaryotic plankton (picoplankton) seemed to fit better the neutral model and possess a larger disperse ability than the larger-sized fraction (nano- and micro-plankton), suggesting that the smaller-sized plankton are less environment filtered and more plastic than the larger-sized plankton. In addition, while interspecific interactions were detected, how much they might contribute to shaping the community structure needs to be explored further in future research.

## Material and methods

### Study area and sample information

During a research cruise aboard Yanping 2, August 6-12, 2016, 49 samples were collected at two stations in the northern South China Sea. Station C6 (117.46°E, 22.13°N) was located at the continental shelf (77 m deep) and station C9 (117.99°E, 21.69°N) at the slope (1369 m deep). As shown in Table S1, sampling targeted two depths at C6, surface (SUR: 0-3 m) and deep chlorophyll maximum (DCM: 35 m at C6 and 75 m at C9), and three depths at C9, with an additional depth named bottom (BOT: 150 m). Sampling frequency was two time points (12:00, 24:00) a day for station C6 and four time points a day (12:00, 18:00, 24:00, 06:00) for C9. For each sample, 40-50 L seawater from more than three CTD rosettes (one CTD rosette contain 12 L seawater) was first filtered through 200 μm bolting cloth to remove large plankton. The seawater was then serially filtered through two size fractions (3 μm and 0.22 μm) using 142-mm diameter polycarbonate membranes (Millipore, Billerica, MA, USA) completed within 20 min. Two size fractions were obtained for each sample: the picoplankton (0.22-3 μm) and nanoplankton/microplankton (3-200 μm). Finally, filters were each stored into a 2-ml tube containing 1 mL lysis buffer (0.1 M EDTA and 1% SDS) and snap-frozen on liquid nitrogen onboard and stored at −80 °C back in the laboratory until DNA extraction. Temperature, salinity, and depth were measured using a CTD profiler. A series of nutrients were measured using continuous-flow Technicon AA3 AutoAnalyzer. The detection limits of nitrate + nitrite (NO_3_^−^+NO_2_^−^), nitrite (NO_2_^−^), phosphate (PO_3_^-^), and silicate (SI_3_O_2_-) were 0.03 μmol L^−1^, 0.04 μmol L^−1^, 0.03 μmol L^−1^ , and 0.05 μmol L^−1^, respectively.

### DNA extraction and high-throughput sequencing (HTS)

Membranes with plankton cells were cut into small pieces and incubated at 56 °C with proteinase K for about three days, with the extended incubation period intended to maximize cell lysis. Then recalcitrant cells that remained unbroken at the end of incubation were mechanically homogenized using the Fastprep^®^-24 Sample Preparation System (MP Biomedicals, USA) with bead-beating (~ 3: 1 mixture of 0.5 mm and 0.1 mm diameter ceramic beads) as previously reported [59]. Next, the DNA extraction was then conducted using a CTAB protocol coupled with Zymo DNA Clean & concentration kit (Zymo Research Corp, Irvine, USA) following Yuan et al. (2015). The concentration and the quality of extracted DNA were determined on NanoDrop-2000 (Thermo Scientific, Wilmington, DE, USA). The V4 region of 18S rDNA was amplified using primers 18SV4-F (GGCAAGTCTGGTGCCAG) and 18SV4-R (GACTACGACGGTATCTRATCRTCTTCG) from 30 ng DNA template of each sample [60]. The PCR products were purified using Agencourt AMPure XP beads and eluted in the Elution buffer. The purified amplicons were quality-checked using the Agilent 2100 Bioanalyzer. The validated libraries were used for sequencing on a MiSeq platform (Illumina, San Diego, CA, USA) with more than 20 Mbp data output of each sample.

Initial data processing was accomplished following the BGI procedure [61]. Briefly, the raw data was polished after removing low quality reads with ambiguous bases (N base), low complexity, adapter contaminations or when the average quality score was less than 20. The high-quality paired-end reads were clustered based on overlap by FLASH [62]. After removing singletons as well as chimeras, the OTUs were determined at a 97% sequencing similarity threshold using UPARSE [63]. Finally, the taxonomic assignment of OTUs was carried out against the SILVA 128 reference database using an 80% bootstrap confidence score [64] on the QIIME platform [65]. Based on the initial assignment of OTUs in the SILVA 128 reference database (https://www.arb-silva.de), we rearranged our OTU taxonomic profile following previous reports [5, 66]. The first classification level included eight super-groups: Alveolata, Amoebozoa, Archaeplastida, Excavata, Incertae_Sedis, Opisthokonta, Rhizaria, and Stramenopiles. OTUs that remained unidentified were binned as Unclassified, and if an OTU showed a similarity < 80% to known species, it was binned into the Undetermined. Before multivariate statistical analysis, the abundance data of OTUs were transformed to relative proportions.

### Diversity analysis

Alpha diversity indices of Sobs, Chao, ACE, Shannon, and Simpson were calculated using the Mothur (v1.31.2) software [67]. Rarefaction curves were constructed using R package RColorBrewer to estimate the sufficiency of sequencing depth to cover the samples’ taxon diversity. Heatmap analysis was made through the R package ade4 with the distance algorithm of euclidean and the complete clustering method. Principal Component Analysis (PCA) was performed through the R package ade4 to visualize differences among samples. The diversity variance analysis was conducted using the Mfuzz package (Parameters: c=9, m=0.8) to cluster samples with the same variable patterns.

### Neutral community model and dispersal ability

A neutral community model was used to determine the potential importance of stochastic process on eukaryotic plankton assembly base on the OTU detection frequency and their relative abundances [9, 68]. The model used in this study was based on the neutral theory [69], which was adapted to suit large microbial populations. In this model, the argument R^2^ represents the overall fit to the neutral model with 95% confidence intervals around all fitting statistics. The argument m represents the immigration rate. In this study, two size fractions were separately calculated for the fit of the neutral model. All computations were performed in R (Version 4.0.2).

To estimate dispersal ability, the pairwise shared proportion of OTUs was calculated, and the average shared proportion was considered as a proxy for dispersal [70]. The linear mixed-effect model with sampling depths as a random effect was used for contrasting the overall difference in dispersal values for two size-fraction communities by using the nlme package in R. Furthermore, results for each water depth were provided with additional comparisons (ANOVA).

### Relationships between eukaryotic communities and environmental and spatial variables

The Variation Partitioning Analysis (VPA) was run in R to quantify the relative effects of environmental and spatial factors in shaping the eukaryotic community, as previously reported [7, 21]. Briefly, before VPA analyses, all environmental variables were z-score transformed to improve normality and homoscedasticity. The spatial variables were then generated using principal coordinates of neighbor matrices (PCNM) analysis based on the latitude and longitude of the sampling sites [71]. Next, forward selection procedures were used to select subsets of environmental variables using the “*ordiR2step*” function from the vegan package in R based on redundancy analysis [72]. Lastly, VPA was conducted using PCNM spatial variables and environmental variables, and ANOVA permutation tests were calculated on the variation explained by each set without the effect of the other. In this analysis, the residual fraction equals the unexplained variance.

After the above forward selection analysis, only temperature and salinity remained potentially influential factors for the entire community structure and used for downstream analyses. For this reason, we analyzed temperature and salinity separately from the other factors. To compare the variance partitioning results, Mantel and partial Mantel tests were run in R using the “vegan” package to determine the correlations between community similarity and environmental and spatial variables.

### Statistical analysis

An analysis of similarity (ANOSIM) was carried out to statistically test the significance of differences in eukaryotic communities between different stations, sampling depths, and diel time points. The argument R in ANOSIM ranges from 0 to 1 represents separation degree between groups and range from −1 to 0 represents separation degree within groups; R=0 indicates no separation, whereas R=1 suggests complete separation. The P-value < 0.05 was taken as statistically significant. Furthermore, the nonparametric Mann-Whitney *U* test and Kruskal-Wallis *H* test from the ‘stats’ package were used to test significant differences of higher eukaryotic taxa among two sampling stations, two sampling times, and three sampling depths.

When less than three groups were compared, the alpha diversity index difference between groups was tested using the Mann-Whitney *U* test. Mantel tests were also conducted to determine correlations between the plankton groups and environmental factors using the “*ggcor*” package. A positive correlation could represent symbiosis, commensalism or parasitism, and a negative correlation means competition or predation.

### Accession number

All the Illumina sequences reported in our study have been submitted to the European Nucleotide Archive (ENA) databank (https://www.ebi.ac.uk/ena/browser/home) under the accession number PRJEB24057 and CNGB Sequence Archive (CNSA) of China National GeneBank DataBase (CNGBdb) (https://db.cngb.org/cnsa/) under the accession number CNP0001483.

## Author contribution

Tangcheng Li and Cong Wang performed experiments. Tangcheng Li, Guilin Liu, Huatao Yuan, Xin Lin, Liying Yu, Ling Li, Yunyun Zhuang and Senjie Lin analyzed the diversity data. Tangcheng Li and Senjie Lin wrote the manuscript.

## Acknowledgements

We wish to thank Bangqin Huang, Xin Liu, Yanping Zhong, and Chentao Guo of Xiamen university for help with sampling and logistic support. We are also indebted to the crew and participants of the Yanping 2 research cruise for assistance in sampling. This work was supported by National Key Research and Development Program of China grant 2016YFA0601202 and the Marine S&T Fund of Shandong Province for Pilot National Laboratory for Marine Science and Technology (Qingdao) (No.2018SDKJ0406-3).

